# Estrogen Deprivation and Periodontitis Interact Across Multiple Tissues

**DOI:** 10.64898/2026.05.15.725533

**Authors:** Nil Yakar, Hatice Hasturk, Carla Alvarez Rivas, Phrao Zimmerman, Zeliha Guney, Birtan Tolga Yilmaz, Yasin Uzun, Philip Trackman, Alpdogan Kantarci

## Abstract

The study investigated the interaction between estrogen deprivation and periodontitis, systemically, in the bone marrow, and locally in periodontal tissues using a mouse model.

**Methods:** We used the ligature-induced periodontitis (LIP) model concurrently with ovariectomy-induced estrogen deprivation. Bone marrow was assessed for myeloid cell proportion by flow cytometry. The femur metaphysis was examined histologically and by micro-CT. Cytokine responses of CD11b^+^ myeloid cells to lipopolysaccharide stimulation were investigated *ex vivo* across ovary-intact (Sham), ovariectomized (OVX), and estrogen-replaced (OVX+E2) mice with or without periodontitis. Estrogen-related alterations in periodontitis, including microbiome composition and transcriptomic changes in the gingiva and dentoalveolar complex, were investigated by 16S rRNA sequencing and bulk RNA sequencing, respectively.

**Results:** Ovariectomy increased osteoblast-like and adipocyte-like cell numbers in femoral marrow, whereas LIP reduced both populations (p = 0.020 and p = 0.029, respectively). LIP increased the bone marrow CD45^+^ hematopoietic fraction in Sham mice. LPS-stimulated bone marrow CD11b^+^ cells from OVX mice showed lower *Tnfα, Ccl2*, and *Il10* expression than Sham mice (p = 0.003, p = 0.005, and p = 0.001, respectively). OVX exacerbated LIP-associated alveolar bone loss, reducing BV/TV (p = 0.003) and increasing osteoclast numbers (p = 0.012). Neither OVX nor E2 replacement significantly altered ligature-associated microbial composition in 16S rRNA sequencing. Bulk RNA sequencing demonstrated estrogen-responsive transcriptomic changes in both the gingiva and dentoalveolar complex, including OVX-associated gene-expression changes that returned toward Sham levels in OVX+E2 mice. These included genes related to stromal regulation (*Acan, Igfbp3, Erbb3*) and immunity (*Gp2, Spib, B2m*).

**Conclusion:** Periodontitis and estrogen deprivation exert combined effects on the bone marrow niche. Estrogen deprivation modulates immune- and healing-related gene expression in the gingiva and remaining dentoalveolar tissues during periodontitis.

---

Periodontitis is a dysbiosis-driven chronic inflammatory disease that has been mechanistically linked to multiple systemic conditions, in part through modulation of the bone marrow microenvironment (Hajishengallis 2022; Hajishengallis and Chavakis 2021). Periodontitis also has been associated with skeletal comorbidities relevant to menopause, particularly osteoporosis (Anbinder et al. 2016; Yu and Wang 2022).

Menopause is characterized by a sharp decline in circulating estrogen levels and represents an inflection point for many health conditions, periodontitis being among them (Davis et al. 2015; Martin et al. 2026). Periodontitis is more prevalent and severe in postmenopausal women, a phenotype often attributed to altered bone turnover (Goyal et al. 2017; Penoni et al. 2017). Importantly, periodontal tissues are hormone-responsive, expressing estrogen receptors in gingiva and periodontal ligament cells (Cao et al. 2007; Quast et al. 2021), as reflected by inflammatory changes during puberty and pregnancy (Guncu et al. 2005). Estrogen deprivation may contribute to periodontitis pathogenesis through reduced estrogen receptor signaling. Although observational studies have suggested a bidirectional link between periodontitis and estrogen deprivation, the local and systemic components of this interaction remain undefined.

We hypothesized that periodontitis and estrogen deficiency interact within the bone marrow microenvironment, where periodontitis worsens skeletal bone loss associated with estrogen deficiency, and within periodontal tissues, where estrogen deficiency exacerbates periodontitis by altering the local inflammatory response. To test this, we first examined whether ligature-induced periodontitis affects bone marrow composition and systemic bone morphometry in ovariectomized mice. We then assessed the impact of estrogen deprivation on ligature-induced alveolar bone loss, microbiome composition, and the inflammatory transcriptional landscape in the gingiva and remaining dentoalveolar complex.

## MATERIALS AND METHODS

A list of key reagents and kits used across the study is provided in Appendix Table 1.

### Animals and procedures

We employed mouse models of ovariectomy-induced estrogen deprivation combined with ligature-induced periodontitis. The study protocol was approved by the ADA Forsyth Institute Animal Care and Use Committee (Protocol #23.009). The experimental unit was one mouse. Female C57BL/6J mice were obtained from The Jackson Laboratory, acclimatized for 1 week, and housed under standard conditions with *ad libitum* access to food and water; mice were 11 weeks old at the start of the experiments. No formal randomization was used for group allocation, and no selection was applied based on body weight or other baseline characteristics. Group allocation and experimental conduct were not blinded; histologic and micro-CT assessments were performed blinded to group identity by coding images and samples before analysis.

Groups included ovary-intact mice (sham-operated, Sham) and ovariectomized mice (OVX), each further divided into periodontitis (Perio) and non-periodontitis (NP) subgroups. Multiple independent cohorts with different timelines were used (Figs. 1a, 2a, and 3c). One cohort also included estrogen-replaced ovariectomized mice (OVX+E2). In this cohort, OVX+E2 mice received subcutaneous estradiol benzoate (1 µg/day in 100 µL corn oil) from day 4 to day 14, whereas Sham and OVX mice received vehicle (Fig. 2a). Endpoint-specific sample sizes are shown as individual data points in the figure panels. Sample size for the 5-week experiment (Fig. 1a) was determined by an a priori power calculation based on systemic bone-density outcomes, whereas sample sizes for the other experiments (Figs. 2a and 3c) were based on experience from this initial experiment. OVX efficiency was confirmed by body-weight monitoring or uterine weight at euthanasia (Appendix Fig. 1a). Ligature-induced periodontitis was established by placing a 5-0 silk suture around the bilateral maxillary second molars and securing it with a palatal knot, as previously described (Kantarci et al. 2020).

**Figure 1.**
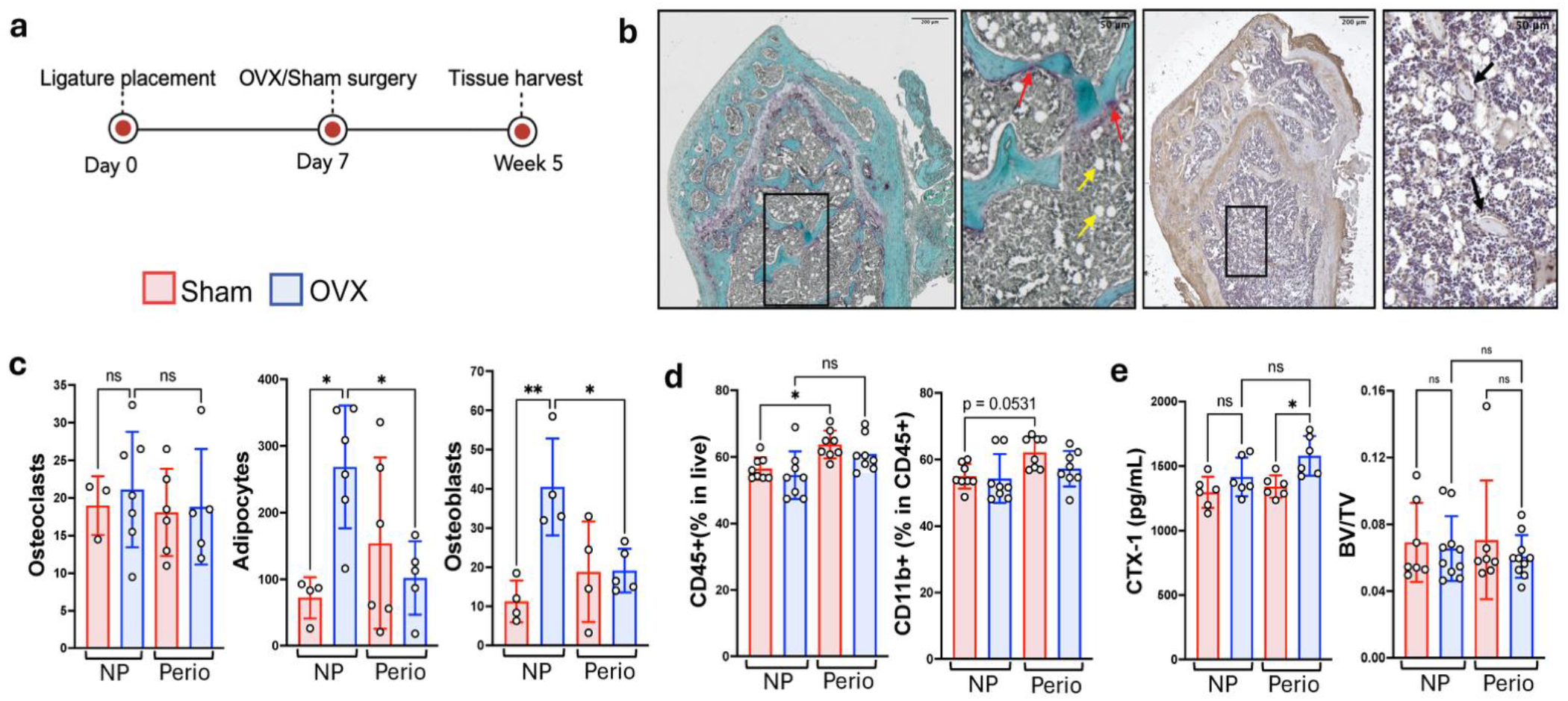
Effects of ovariectomy and periodontitis on cellular composition of bone marrow and bone. **a**, Experimental timeline. b, Representative TRAP-stained and osteocalcin-immunostained distal femur sections. Red arrows indicate TRAP-positive osteoclast-like cells, yellow arrows indicate adipocyte-like cells, and black arrows indicate osteocalcin-positive osteoblast-like cells. c, Bar graphs of osteoclast, adipocyte, and osteoblast counts in the secondary spongiosa. d, Percentages of CD45^+^ and CD11b^+^ cells in bone marrow assessed by flow cytometry. e, Serum C-terminal telopeptide of type I collagen (CTX-1) levels measured by ELISA and femoral trabecular bone volume fraction (BV/TV) measured by micro-CT. Sham: sham-operated; OVX: ovariectomized; NP: non-periodontitis; Perio: periodontitis. Asterisks indicate statistical significance: p < 0.05; p < 0.01; ns: not significant.

**Figure 2.**
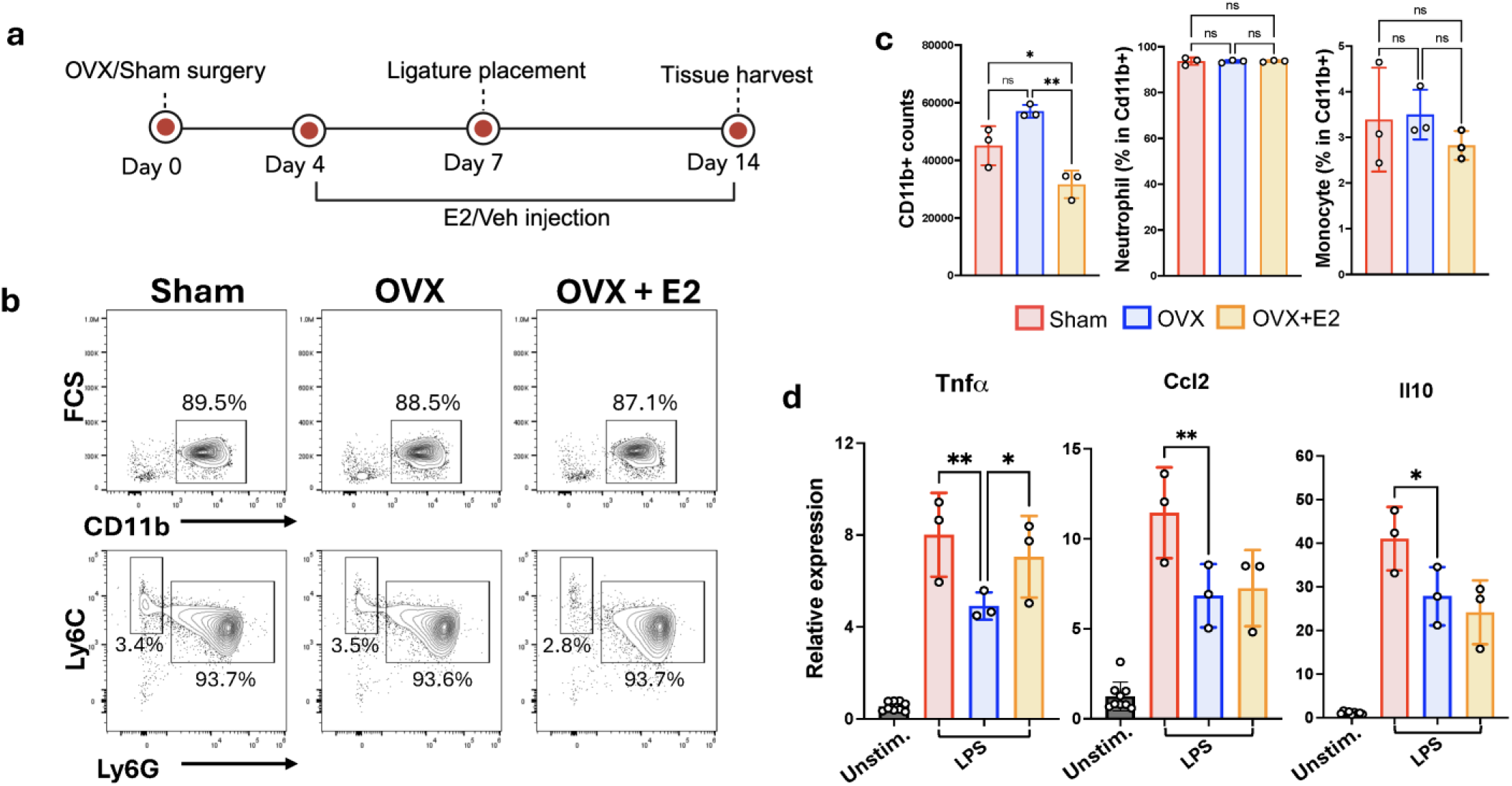
LPS-induced cytokine expression in CD11b^+^-enriched bone marrow cells. a, Experimental timeline; all three groups underwent ligature-induced periodontitis. b, Representative flow-cytometry gating strategy. Neutrophils were defined as CD11b^+^Ly6G^+^Ly6C^lo/int^ cells, and monocytes were defined as CD11b^+^Ly6G^−^Ly6Chi cells. c, Quantification of live CD11b^+^ cell yield and proportions of neutrophils and monocytes within the CD11b^+^ population across groups. d, Relative expression of LPS-induced Tnfα, Ccl2, and Il10 in bone marrow CD11b^+^ cells. Unstimulated CD11b^+^ cells served as controls. Sham: sham-operated, OVX: ovariectomized, OVX+E2: ovariectomized with estrogen replacement. Asterisks indicate statistical significance: *p < 0.05; **p < 0.01; ns: not significant.

### Histology

Femurs and maxillary dentoalveolar complexes were decalcified in 10% EDTA (pH 7.4) for 14 and 21 days, respectively, then embedded in paraffin and sectioned at 5 µm.

TRAP staining was performed using a Leukocyte Acid Phosphatase kit according to the manufacturer’s instructions. Slides were counterstained with Fast Green. Brightfield images were acquired using a Zeiss Axio Observer Z1/7 microscope equipped with an Axiocam 712 camera and a 20×/0.8 air objective. TRAP-positive cells with osteoclast-like morphology located along bone surfaces were quantified in the secondary spongiosa of the distal femoral metaphysis and between the first and third molars in the alveolar bone region.

Immunohistochemistry was performed for osteocalcin staining as detailed in the Appendix. Osteocalcin-positive cells lining trabecular bone surfaces were quantified as osteoblast-like cells in the secondary spongiosa of the distal femoral metaphysis. Bone marrow adipocytes were quantified within the same region based on morphological appearance as large unilocular vacuolated cells.

### Bone marrow flow cytometry assay

Bone marrow cells were harvested from femurs and tibiae by flushing with PBS, followed by red blood cell lysis. For bone marrow flow cytometry analysis, cells were stained with a viability dye and antibodies against CD45, CD11b, Ly6G, and Ly6C. Data were acquired on an Attune NxT flow cytometer (Thermo Fisher) and analyzed using FlowJo software (v10). Additional details on flow cytometry assay are provided in the Appendix.

### Bone resorption and morphometric analyses

Serum C-terminal telopeptide of type I collagen (CTX-1), a marker of bone resorption, was quantified by ELISA.

Femurs and maxillae were scanned by micro-CT (Scanco Medical, µCT40). Three-dimensional reconstruction and morphometric analyses were performed using Scanco software (v5.0). Trabecular bone parameters, including bone volume fraction (BV/TV), trabecular number (Tb.N), trabecular thickness (Tb.Th), and trabecular separation (Tb.Sp), were calculated for the distal femoral metaphysis and the maxillary second molar alveolar bone region. Scan settings and region-of-interest definitions are provided in the Appendix.

### *Ex vivo* stimulation and RT-qPCR of bone marrow CD11b^+^ cells

Bone marrow cells were flushed from femurs and tibiae using PBS, and red blood cells were lysed. CD11b^+^ cells were enriched using magnetic microbeads. Cells were stimulated with 100 ng/mL *E. coli* LPS for 4 hours. A quarter of the enriched cell population from each sample was analyzed by flow cytometry to assess pre-LPS cell composition using antibodies against CD11b, Ly6G, and Ly6C. Within the enriched CD11b^+^ population, neutrophils were gated as CD11b^+^Ly6G^+^Ly6C^lo/int^ cells, and monocytes were gated as CD11b^+^Ly6G^−^Ly6C^hi^ cells. Flow cytometry data acquisition and analysis were performed as described above. Representative gating strategies are shown in Fig. 2b. At the end of LPS stimulation, cells were lysed in RLT buffer. RNA was extracted and reverse transcribed. RT-qPCR was performed for Actb, Tnfα, Ccl2 and Il10. Relative gene expression levels were calculated against the unstimulated controls group using the ΔΔCt method.

### 16S rRNA sequencing

Retrieved silk ligatures were placed in Pras Dilution Blank, and DNA was extracted using the MasterPure Complete DNA Purification Kit. Libraries targeting the V1–V3 region were prepared and sequenced by Zymo Research Corp. (CA, USA) on an Illumina NextSeq 2000 platform. Downstream sequence processing, taxonomic assignment, and diversity analyses were performed by the Forsyth Oral Microbiome Core as detailed in the Appendix.

### Bulk-RNA sequencing

Gingiva and the dentoalveolar complex of the maxillary molar region were flash-frozen and pulverized under liquid nitrogen. Tissues were lysed in RLT Plus buffer, homogenized using QIAshredder columns, and RNA was isolated using a column-based kit followed by cleanup. Library preparation and sequencing were performed by Quintara Biosciences (MA, USA), as detailed in the Appendix.

Sequencing reads were aligned to the mouse reference genome GRCm38/mm10, and downstream analyses were performed in R. Differential expression analysis was performed using DESeq2. Adjusted p values were calculated using the Benjamini–Hochberg procedure implemented in DESeq2, and results were summarized using an adjusted p-value threshold of 0.10. DESeq2 independent filtering and outlier handling were applied according to the standard workflow (Love et al. 2014). Pathway enrichment analysis was performed using MSigDB Hallmark gene sets (Liberzon et al. 2015) following the GSEA framework (Subramanian et al. 2005). To visualize sample-level pathway patterns, gene set variation analysis (GSVA) scores were calculated for OVX-altered Hallmark gene sets and displayed as row z-score heatmaps (Appendix Fig.2). OVX-altered and E2-recovered genes and Hallmark gene sets were defined as those differing significantly between OVX and Sham and between OVX and OVX+E2, whereas OVX+E2 either did not differ significantly from Sham or shifted beyond Sham in the opposite direction.

### Statistical Analysis

Statistical analyses were performed using GraphPad Prism (v10). Student’s t-test was used for comparisons between two groups. For three or more groups, ordinary one-way ANOVA followed by Šídák’s multiple-comparisons test was used for pairwise comparisons.

## RESULTS

### Periodontitis affects bone marrow immune and stromal cell composition

To evaluate the impact of ligature-induced periodontitis on bone marrow, periodontitis was initiated 1 week before sham or ovariectomy surgery, and the total experimental period lasted 5 weeks (Fig. 1a). This cohort included four groups: Sham and OVX, each including non-periodontitis (NP) and periodontitis (Perio) subgroups. Representative TRAP-stained and osteocalcin-immunostained distal femur sections are shown in Fig. 1b. TRAP staining in femur did not show a significant difference in osteoclast-like cell numbers. However, secondary spongiosa adipocyte and osteoblast-like cell numbers were significantly increased with ovariectomy (p=0.015 and p=0.003, respectively) and both were reduced with periodontitis (p=0.029 and 0.020, respectively) (Fig. 1c).

Flow cytometry assay of the bone marrow demonstrated an increased proportion of total leukocytes (CD45^+^ cells) in periodontitis groups, reaching significance in Sham Perio versus Sham NP (p=0.036). CD11b^+^ Myeloid cells comprised approximately 48-68% of CD45^+^ cells (Fig. 1d). There were no significant differences in the percentage of CD11b^+^Ly6G^+^Ly6C^lo/int^ neutrophils or CD11b^+^Ly6G^-^Ly6C^hi^ monocytes within the CD11b^+^ Myeloid compartment between groups.

We further investigated the impact of periodontitis on systemic bone resorption using serum CTX-1 levels and femoral trabecular bone morphometry. Although serum CTX-1 levels showed an increasing trend with both ovariectomy and periodontitis, micro-CT analysis of the femur metaphysis did not reveal significant differences in bone morphometric parameters including BV/TV, Tb.N, Tb.Th, or Tb.Sp. across groups (Fig. 1e and Appendix Fig. 1c).

### Estrogen deprivation attenuates neutrophil responsiveness to LPS stimulation in the bone marrow during ligature-induced periodontitis

It has been previously established that estrogen modulates myeloid cell development and cytokine responses to TLR4 stimulation (Calippe et al. 2010). To further investigate the impact of estrogen deprivation on myeloid cell responses during periodontal inflammation, we used the ligature induced periodontitis model on Sham, OVX and OVX+E2 groups. Fig. 2a summarizes the experimental timeline. Bone marrow CD11b^+^ cell enrichment yielded a population composed of approximately 88% CD11b^+^ cells; representative gating is shown in Fig. 2b. The number of recovered live CD11b^+^ cells was lower in the OVX+E2 group than in both the Sham and OVX groups (p = 0.048 and p = 0.002, respectively; Fig. 2c). Neutrophils, defined as CD11b^+^Ly6G^+^Ly6C^lo/int^ cells, comprised the majority of the enriched CD11b^+^ fraction (approx. 93%), whereas monocytes, defined as CD11b^+^Ly6G^−^Ly6C^hi^ cells, represented a minor fraction. The frequencies of neutrophils and monocytes were not significantly different between Sham, OVX, and OVX+E2 groups (Fig. 2c).

After 4 hours of *in vitro* LPS stimulation of the bone marrow CD11b^+^-enriched cells, RT-qPCR was performed to assess Tnfα, Ccl2, and Il10 mRNA expression. LPS stimulation increased the mRNA levels of all three genes relative to unstimulated controls. Tnfα levels were significantly lower in OVX than in Sham and OVX+E2 groups (p=0.003 and p=0.039, respectively). Ccl2 and Il10 levels were likewise lower in OVX than in Sham (p=0.005 and p=0.001, respectively) (Fig. 2d).

### Estrogen deprivation exacerbates ligature-induced alveolar bone loss without altering the ligature-associated microbiome

Alveolar bone morphometry around maxillary second molar was analyzed by micro-CT, which demonstrated lower BV/TV in the ovariectomized mice with periodontitis (OVX Perio) compared with Sham Perio (p=0.003) (Fig. 3a). Greater alveolar bone loss in the ovariectomized group was also supported by trends in Tb.Th and Tb.Sp (Appendix Fig. 1d). Osteoclast numbers were also higher in OVX Perio compared with Sham Perio (p=0.012) (Fig. 3b).

**Figure 3.**
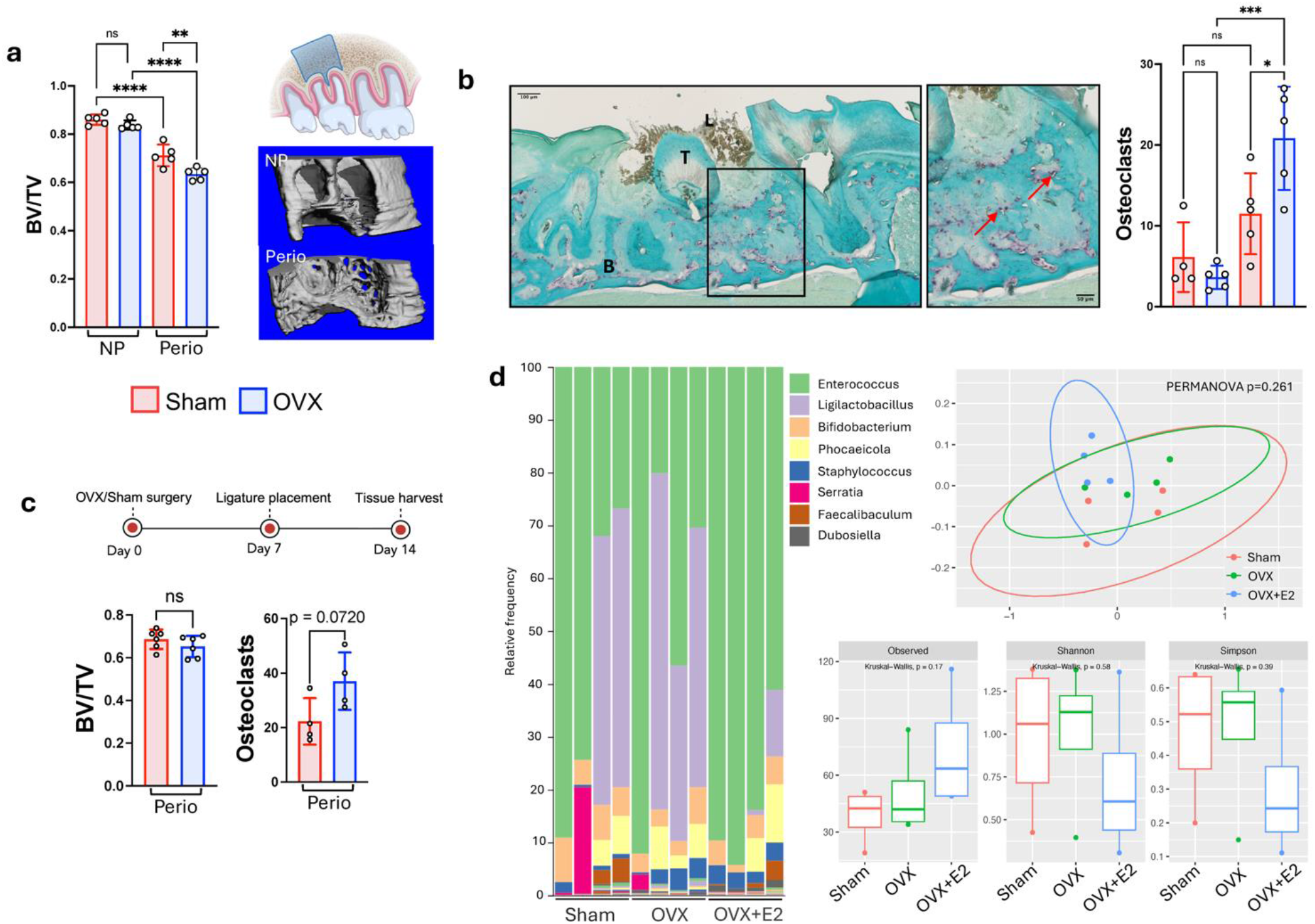
Alveolar bone phenotypes and ligature-associated microbiome. **a**, Bar graph of alveolar bone BV/TV across groups, with representative reconstructed micro-CT images of the alveolar bone region surrounding the maxillary second molar. The experimental timeline is the same as in Figure 1a. **b**, Representative TRAP-stained alveolar bone sections (B, bone; T, tooth; L, ligature) and the bar graph shows osteoclast-like cell counts in the interdental region between the first and third molars. The experimental timeline is the same as in Figure 1a. **c**, Shorter experimental timeline and bar graphs of alveolar bone BV/TV and osteoclast-like cell counts in Sham Perio and OVX Perio mice. **d**, Relative abundance of the eight most abundant genera recovered from silk ligatures and alpha-diversity metrics (observed richness, Shannon index, and Simpson index). The experimental timeline and groups are the same as in Figure 2a. Principal coordinates analysis (PCoA) of Bray–Curtis dissimilarity is shown at the genus level. Sham: sham-operated; OVX: ovariectomized; NP: non-periodontitis; Perio: periodontitis. Asterisks indicate statistical significance: p < 0.05; p < 0.01; p < 0.001.

Reportedly, ligature-induced periodontitis induces alveolar bone loss in 1 week (Abe and Hajishengallis 2013), whereas OVX induces systemic skeletal changes after several weeks (Sophocleous and Idris 2014). We tested a shorter experimental timeline to investigate earlier effects of estrogen deprivation on the inflammatory bone loss, while minimizing the impact of systemic osteoporosis. In this model, we compared Sham Perio and OVX Perio groups, in which surgery was performed on day 0, followed by periodontitis induction on day 7 and euthanasia on day 14 (Fig. 3c). Micro-CT analyses of alveolar bone did not show significant differences between groups, whereas TRAP-positive osteoclast-like cells showed a trend toward higher levels in OVX Perio (p=0.072) (Fig. 3c). We used this experimental design to investigate the impact of estrogen deprivation on the ligature-associated microbiome by 16S rRNA sequencing in Sham, OVX, and OVX+E2 groups. No significant differences in microbial composition were detected between the three groups, based on alpha- and beta-diversity metrics and relative microbial abundances (Fig. 3d). The only exception was the bacterium *Mammaliicoccus lentus*, which was more abundant in Sham, compared to OVX (W = 41 of 137 tested taxa).

### Estrogen deprivation alters the transcriptome in the dentoalveolar complex and gingiva

To assess estrogen-responsive gene expression within the dentoalveolar complex and gingiva during periodontitis, we performed bulk RNA sequencing of samples from Sham, OVX, and OVX+E2 mice, all of which received ligature applications. Differentially expressed genes were identified in both tissues using an adjusted p-value threshold of 0.10 without a fold-change threshold, as visualized in the volcano plots shown in Fig. 4a, b.

**Figure 4.**
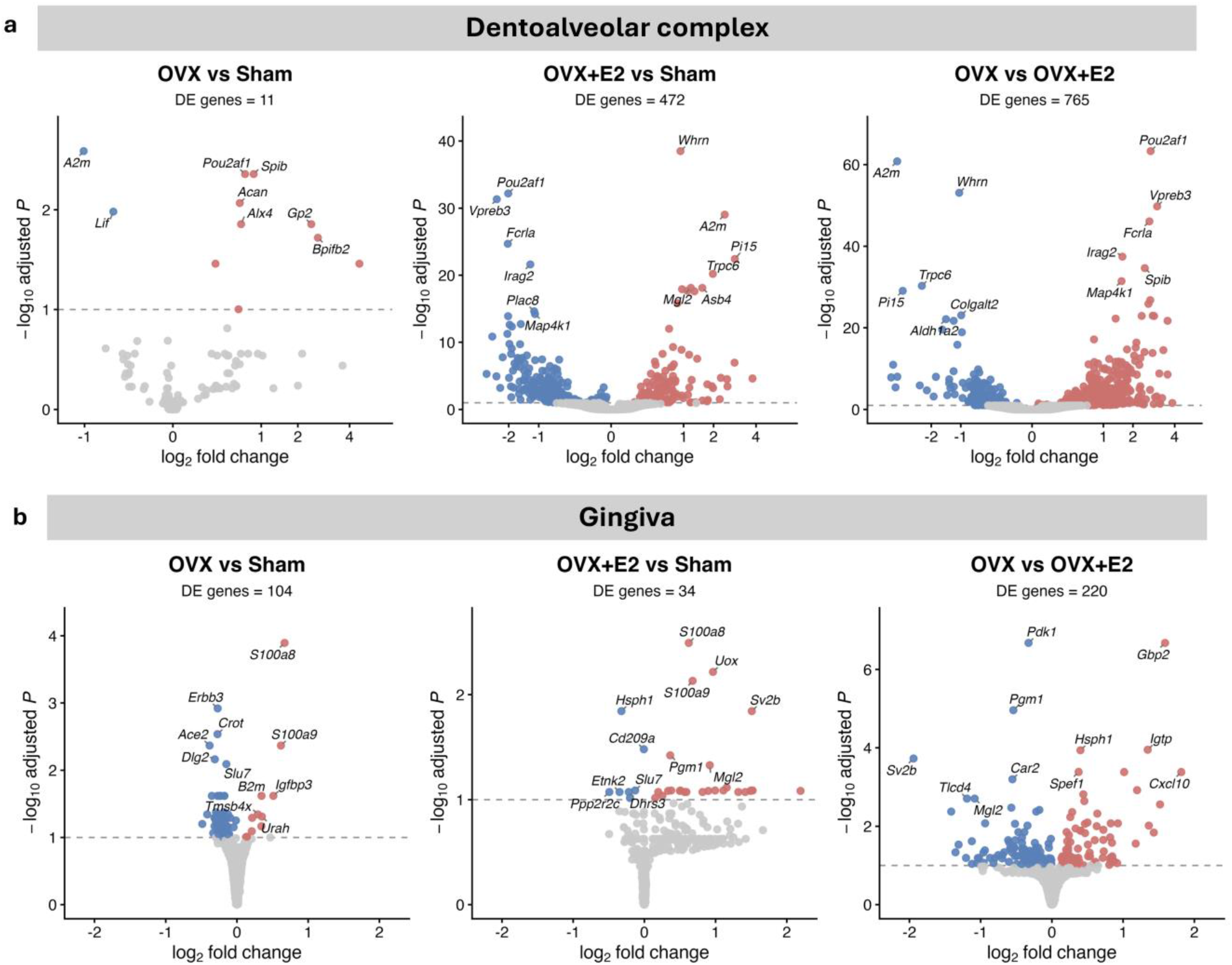
RNA sequencing in the dentoalveolar complex and gingiva. Experimental timeline is the same as in Figure 2a. All groups underwent ligature-induced periodontitis. **a**, Volcano plots of differentially expressed genes in pairwise comparisons in the dentoalveolar complex (*padj* < 0.1). **b**, Volcano plots of differentially expressed genes in pairwise comparisons in the gingiva (*padj* < 0.1). Sham: sham-operated; OVX: ovariectomized; OVX+E2: ovariectomized with estrogen replacement.

Within the dentoalveolar complex, 11 genes were differentially expressed in Sham vs. OVX (9 up, 2 down), 472 genes in OVX+E2 vs. Sham (210 up, 262 down), and 765 genes in OVX vs. OVX+E2 (447 up, 318 down). These findings indicate that OVX-associated transcriptomic changes in the dentoalveolar complex were relatively modest at this time point, whereas estrogen injection induced a broader transcriptional shift (Fig. 4a). In gingiva, 104 genes were differentially expressed in Sham vs. OVX (11 up, 93 down), 34 genes in OVX+E2 vs. Sham (27 up, 7 down), and 220 genes in OVX vs. OVX+E2 (92 up, 128 down), indicating diverse ovariectomy-associated transcriptional response in gingiva at this time point (Fig. 4b).

Ovariectomy-associated differentially expressed genes that were normalized or overexpressed after estrogen supplementation are summarized in Figure 5. In the dentoalveolar complex, eight OVX-associated genes showed E2-associated normalization or over-recovery (padj < 0.1). Genes with lower abundance in OVX included A2m and Lif, whereas genes with higher abundance included *Acan, Alx4, Fcrla, Gp2, Pou2af1*, and *Spib*, spanning matrix remodeling and immunity-associated programs (Fig. 5a). In gingiva, ten ovariectomy-associated genes showed E2-associated normalization. Genes with lower abundance in OVX included *Atp9a, Chil4, Erbb3, Il1rl2, Marchf7*, and *Pdk1*, whereas genes with higher abundance included *B2m, Hnrnpc, Hspe1*, and *Igfbp3*. These gene sets indicate involvement of pathways spanning repair and immune response (Fig. 5b).

**Figure 5.**
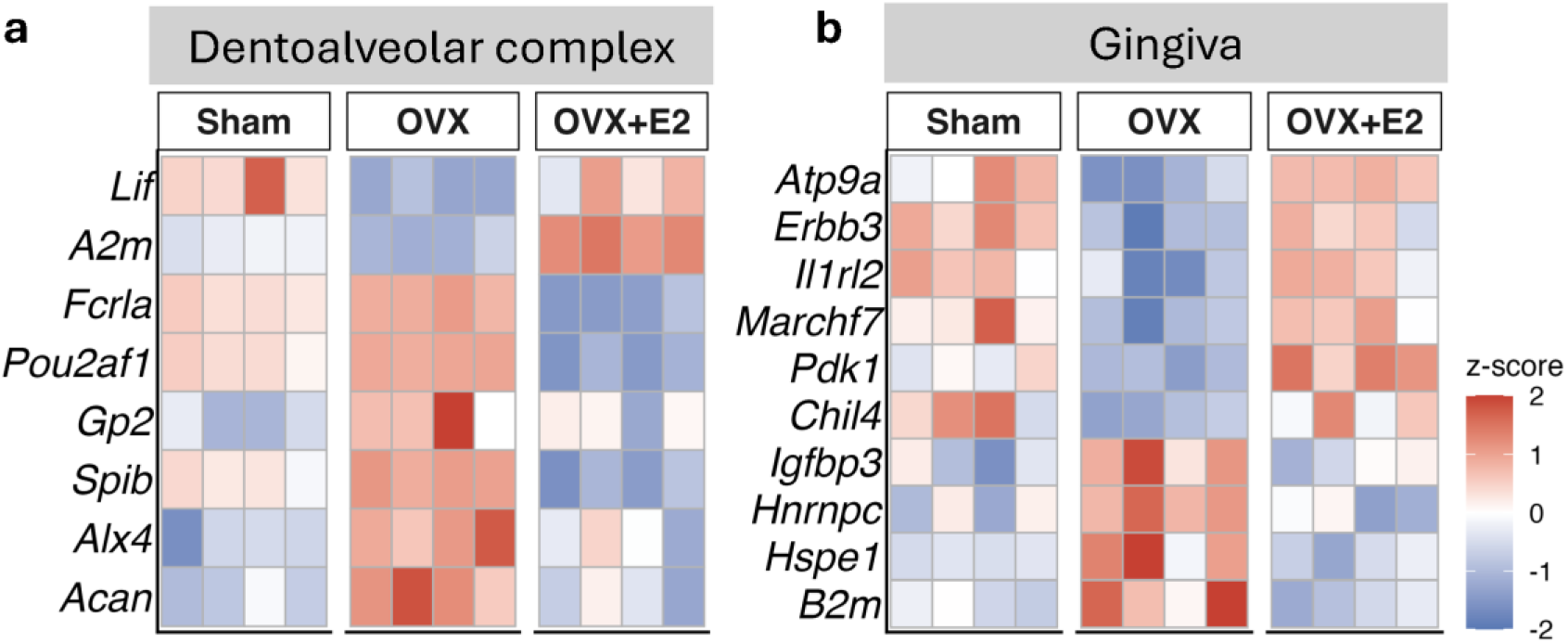
Heatmaps of ovariectomy-associated genes showing E2-responsive patterns. Experimental timeline is the same as in Figure 2a. All groups underwent ligature-induced periodontitis. **a**, Heatmap of ovariectomy-associated differentially expressed genes in the dentoalveolar complex that were normalized or over-recovered by estrogen replacement. **b**, Heatmap of ovariectomy-associated differentially expressed genes in the gingiva that were normalized by estrogen replacement. Sham: sham-operated; OVX: ovariectomized; OVX+E2: ovariectomized with estrogen replacement.

We next performed pre-ranked gene set enrichment analysis (GSEA) using MSigDB Hallmark gene sets to explore pathway-level changes associated with OVX and E2 treatment. Appendix Figure 2 summarizes gene sets that were reduced or increased by OVX and then shifted toward, or beyond, the Sham pattern after E2 injection. In the dentoalveolar complex, OVX reduced gene sets related to tissue organization and energy metabolism, including apical junction, epithelial-mesenchymal transition, and glycolysis, while increasing interferon-response and cell-cycle gene sets. In gingiva, OVX reduced protein secretion and metabolic gene sets and increased inflammatory gene sets, including inflammatory response and TNFα signaling via NF-κB. Together, these findings suggest that estrogen deprivation shifts periodontal tissues away from tissue organization and metabolic programs and toward immune activation (Appendix Fig. 2a, b).

## DISCUSSION

In this study, we provide experimental evidence that supports the bidirectional interactions between estrogen deficiency and periodontitis. Estrogen deprivation modifies systemic and local responses to periodontal inflammation. Both ovariectomy-induced estrogen deficiency and periodontitis modulated the femoral bone marrow cellular composition, while ovariectomy exacerbated the inflammatory alveolar bone loss, altered the *ex vivo* LPS response of bone marrow-derived CD11b^+^ neutrophils, and altered estrogen-responsive transcriptional programs in the gingiva and dentoalveolar complex.

Periodontitis was associated with reduced adipocyte- and osteoblast-like cell numbers in the bone marrow of ovariectomized mice and with an increased frequency of CD45^+^ bone marrow cells in ovary-intact mice. Previous evidence has demonstrated that the inflammatory stress during periodontitis can drive emergency myelopoiesis and remodel the marrow microenvironment (Li et al. 2022; Mitroulis et al. 2020). Since estrogen deprivation also affects the bone marrow (Kitajima et al. 2015), periodontitis-driven marrow changes may contribute to menopause-related systemic comorbidities, including osteoporosis. We tested this question by assessing the systemic bone resorption and femoral trabecular bone morphometry in our combined model. Although serum CTX-1 showed directional increases with ovariectomy and periodontitis, femoral micro-CT did not support a cumulative effect of periodontitis on trabecular bone loss in this model. A limitation of our model is the relatively low peak bone mass of C57BL/6 mice and the variable skeletal response to ovariectomy in this strain which may in part explain the lack of significant difference (Butylina et al. 2025).

We next examined whether estrogen deprivation alters the inflammatory response of bone marrow CD11b^+^ cells during periodontitis. The enriched CD11b^+^ fraction was predominantly composed of neutrophils, which are estrogen-responsive cells (Molero et al. 2002). LPS-stimulated CD11b^+^ cells from ovariectomized mice showed reduced expression of *Tnfα, Ccl2*, and *Il10* compared with Sham mice, with *Tnfα* restored by E2 replacement. These findings suggest that estrogen deprivation may be linked to a tolerance-like state in bone marrow neutrophils after bacterial stimulation.

Ovariectomy exacerbated ligature-induced alveolar bone loss and increased osteoclast-like cell numbers. In line with this, epidemiological evidence links menopause to greater periodontal attachment loss, often attributed to low bone mineral density (Qi et al. 2023). Other contributors, including the microbiome and immune response, have been proposed, but experimental data remain limited (Martin et al. 2026). In our model, estrogen deprivation did not produce a major shift in ligature-associated microbial composition. This finding is consistent with our previous clinical observation that postmenopausal women had greater attachment loss without a dysbiotic shift in the subgingival microbiome (Yakar et al. 2025). We therefore examined host transcriptional changes in the gingiva and dentoalveolar complex.

In the dentoalveolar complex, ovariectomy-associated genes included matrix and skeletal-development genes, such as *Acan* and *Alx4* (Dateki 2017; Lan et al. 2024). Other differentially expressed genes were linked to immunity, including *Gp2, Fcrla, Pou2af1*, and *Spib* (Hase et al. 2009; Reshetnikova et al. 2012; Sasaki et al. 2012; Zhao et al. 2008). These findings suggest that estrogen deprivation affects both matrix-remodeling and immune programs within the dentoalveolar compartment.

In the gingiva, ovariectomy-responsive genes included repair-linked genes with lower abundance and stromal and immune-regulatory genes with higher abundance. Genes with lower abundance in ovariectomy included *Erbb3, Pdk1, Atp9a, Chil4*, and *Il1rl2*, which are linked to epithelial repair, cell migration, vesicle trafficking, and mucosal healing (Gagliardi et al. 2015; Scheibe et al. 2017; Wang et al. 2021; Zhang et al. 2012). *Igfbp3* was increased with ovariectomy. IGFBP-3, insulin-like growth factor-binding protein 3, is a secreted protein that regulates insulin-like growth factor (IGF-1) availability and signaling. Human gingival fibroblasts can release both insulin-like growth factor 1 (IGF-1) and IGFBP-3; thus, its expression shift may indicate a fibroblast-linked regulatory axis in the estrogen-deprivation-related periodontitis phenotype (Saygun et al. 2008; Varma Shrivastav et al. 2020).

The MSigDB Hallmark analysis supported our biological interpretation of differentially expressed genes. Ovariectomy reduced pathways associated with tissue organization, protein secretion, and metabolism, while increasing inflammatory and immune-response pathways, particularly in the gingiva. Taken together, the differentially expressed genes and pathway data suggest that estrogen deprivation shifts inflamed periodontal tissues away from repair and metabolic activity and toward immune activation.

Despite the coherence of our observations, we acknowledge the study limitations. First, the dentoalveolar complex was analyzed as a composite tissue containing alveolar bone, periodontal ligament, bone marrow, and tooth, which limits compartment-specific interpretation. Second, the number of E2-responsive genes was modest, likely due to the low abundance of estrogen-responsive cells in periodontal tissues; nevertheless, the concordance between gene-level and Hallmark pathway patterns supports a consistent estrogen-related signal. Estrogen receptor-enriched methods may define estrogen-responsive mechanisms in greater resolution.

In conclusion, our findings support our hypothesis that estrogen deprivation modulates periodontitis not only through altered bone turnover, but also through effects on bone marrow immune cells, gingival repair and immune programs. These observations broaden the current view of postmenopausal periodontitis and support further investigation of estrogen-responsive mechanisms in periodontal tissues.

## Conflict of Interest Statement

The authors declare no conflicts of interest with respect to the authorship and publication of this article.

### Acknowledgements

The authors would like to thank Yaser Peymanfar for guidance during the bone analyses and Dorathy Vargas for help with the experimental work. This work was supported by ADA Forsyth Pilot Grants (AFIPLT01), NIA RF1AG062496 (to AK), NIDCR R90DE027638 (to NY), and TUBITAK 2214/A (to NY) and 2219 (to ZG). ADA Forsyth Micro-CT, Oral Microbiome, Flow Cytometry, and Microscopy Core facilities were used for data acquisition. Illustrations were generated in BioRender. ChatGPT 5.4 (OpenAI) was used for language refinement.

## Author contributions

NY, CAR, and AK designed the study. NY, PZ, and ZG performed the experiments. NY and BTY performed tissue processing and downstream analyses. YU assisted with data analysis. NY wrote the manuscript. HH, PT and AK supervised the study and critically reviewed the manuscript. All authors gave their final approval and agree to be accountable for all aspects of the work.

## Data Availability

Raw bulk RNA-sequencing data have been deposited in NCBI GEO under accession number GSE327719. Raw 16S rRNA sequencing data have been deposited in NCBI SRA under accession number PRJNA1456169.

