## Supplementary Appendix for "Estrogen Deprivation and Periodontitis Interact Across Multiple Tissues"

**Appendix Table 1: Key resources**

| **Reagents or kits** | **Source** | **Identifier** |
|---|---|---|
| **Injections / Histology/ Bone resorption** | | |
| Estradiol benzoate | Sigma-Aldrich | #E8515 |
| Corn oil | Sigma-Aldrich | #C8267 |
| Leukocyte Acid Phosphatase kit | Sigma-Aldrich | #387A |
| Fast Green FCF | Sigma-Aldrich | #F7252 |
| Anti-osteocalcin antibody | Abcam | #ab93876 |
| Goat anti-rabbit IgG antibody | Abcam | #ab6721 |
| Rabbit IgG isotype control antibody | Invitrogen | #02-610-2 |
| DAB substrate kit | Cell Signaling Technology | #8059 |
| CTX-1 ELISA kit | MyBioSource | #MBS8801224 |
| **Bone marrow flow cytometry** | | |
| Live/Dead Yellow viability dye | Invitrogen | #L34959 |
| anti-CD45-PE | BioLegend | B315677 |
| anti-CD11b-PerCP/Cy5.5 | BioLegend | B375903 |
| anti-Ly6G-BV711 | BioLegend | #127643 |
| anti-Ly6C-BV421 | BioLegend | B391725 |
| ABC Total Antibody Compensation Bead Kit | Invitrogen | A10497 |
| ***Ex vivo* LPS-stimulation experiment** | | |
| CD11b MicroBeads | Miltenyi | #130-097-142 |
| LS columns | Miltenyi | #130-042-401 |
| *E. coli* LPS O111:B4 | Sigma-Aldrich | #L2630 |
| Live/Dead Yellow viability dye | Invitrogen | #L34959 |
| anti-CD11b-APC | BioLegend | B481850 |
| anti-Ly6G-PE | BioLegend | B476627 |
| anti-Ly6C-FITC | BioLegend | B477805 |
| ABC Total Antibody Compensation Bead Kit | Invitrogen | A10497 |
| RLT buffer | Qiagen | #79216 |
| SuperScript IV VILO | Invitrogen | #11756050 |
| TaqMan Fast Advanced Master Mix | Applied Biosystems | #4444558 |
| Actb TaqMan probe | Thermo Fisher | Mm02619580_g1 |
| Ccl2 TaqMan probe | Thermo Fisher | Mm00441242_m1 |
| Tnfα TaqMan probe | Thermo Fisher | Mm00443258_m1 |
| Il10 TaqMan probe | Thermo Fisher | Mm01288386_m1 |
| **16S rRNA sequencing** | | |
| Pras Dilution Blank | Anaerobe Systems | #AS-910 |
| MasterPure Complete DNA Purification Kit | Biosearch Technologies | #NC9801456 |
| **Bulk RNA-seq** | | |
| RNeasy Mini Kit | Qiagen | #74104 |
| RLT Plus buffer | Qiagen | #1053393 |

| | | |
|---|---|---|
| QIAshredder columns | Qiagen | #79656 |
| RNA Clean & Concentrator kit | Zymo Research | #R1016 |
| VAHTS Universal V10 RNA-seq Library Prep Kit | Vazyme | #NR616-02 |

**Animals and procedures**

Ovariectomy or sham surgeries were performed under ketamine/xylazine anesthesia (ketamine 87.5 mg/kg and xylazine 12.5 mg/kg, intraperitoneally). The peritoneal cavity was accessed through a midline ventral incision. In OVX groups, both ovaries were removed while preserving the uterus. In sham groups the ovaries were located and left intact. Muscle was closed with chromic gut, and skin was closed with 5-0 polypropylene sutures. Postoperative analgesia was provided with a single dose of buprenorphine (Ethiqa XR, 3.25 mg/kg, subcutaneously).

At the end of the experimental period, mice were euthanized by CO2 asphyxiation. Whole blood was collected from the inferior vena cava and allowed to clot for 1 h at room temperature. Samples were centrifuged at 2,000 × g for 10 min, and serum supernatants were collected and stored at −80 °C until analysis. Femurs and maxillae designated for micro-CT and histology were fixed in 10% formalin overnight and then transferred to 70% ethanol. Femurs and tibiae designated for flow cytometry or CD11b+ cell separation were collected in PBS and processed immediately.

**Histology**

Osteocalcin immunohistochemistry was performed on paraffin sections with heat-mediated antigen retrieval. Sections were incubated overnight at 4 °C with rabbit anti-osteocalcin primary antibody diluted 1:500 in blocking buffer, followed by goat anti-rabbit IgG secondary antibody diluted 1:1000 for 1 h at room temperature. Rabbit IgG isotype control antibody was used for the controls. Signal was developed using DAB, and sections were counterstained with Gill's hematoxylin.

For femoral histomorphometric analyses, the secondary spongiosa was defined in Fiji/ImageJ. The growth plate was manually traced, and parallel reference lines were generated at 250 μm and 1000 μm distal to the growth plate. The trabecular region between these two lines was used as the secondary spongiosa region.

**Bone marrow flow cytometry assay**

Compensation was performed using compensation beads according to the manufacturer's instructions. The gating strategy was as follows: debris was excluded based on forward- and side-scatter, followed by singlet gating. Live cells were identified by exclusion of Live/Dead-positive events. Leukocytes were defined as $CD45^+$ cells, and myeloid cells were identified as $CD11b^+$ cells within the $CD45^+$ population. Within the $CD11b^+$ gate, $Ly6G^+Ly6C^{lo/int}$ cells were defined as neutrophils and $Ly6G^-Ly6C^{hi}$ cells were defined as monocytes.

**Bone morphometric analyses**

Micro-CT scans were acquired at 70 kVp, 114 μA, and 6 μm voxel size. For femur trabecular bone density analyses, the region of interest was defined in the distal metaphysis beginning 100 slices distal to the growth plate and extending for 150 consecutive slices. Trabecular parameters were calculated using a threshold of 132.

For maxillary alveolar bone density analyses, the region of interest was centered on the maxillary second molar. The mesial and distal boundaries of the second molar were identified, and the midpoint between them was used as the center of the region of interest. Fifty consecutive slices on each side of this midpoint were analyzed, corresponding to a total of 100 slices, using a threshold of 184.

**16S rRNA sequencing analysis**

Raw reads were processed using the DADA2 pipeline for quality filtering, denoising, paired-read merging, and chimera removal. Species-level taxonomic assignment was performed using an open-reference BLASTN-based pipeline against a combined reference database including HOMD v15.22, MOMD v5.1, and the NCBI 16S rRNA reference database. Alpha diversity was evaluated at the species level using Observed richness, Shannon diversity, and Simpson diversity indices, with between-group differences assessed by Kruskal–Wallis testing. Beta diversity was assessed using Bray–Curtis dissimilarity, and group differences were tested by PERMANOVA. Pairwise differential abundance analysis was performed using ANCOM-BC2.

**Bulk RNA sequencing and analysis**

RNA quality was assessed using the Agilent TapeStation, and samples with an RNA integrity number (RIN) > 7.0 were used for library preparation. Libraries were prepared using the VAHTS Universal V10 RNA-seq Library Prep Kit with poly(A) mRNA enrichment. Libraries were sequenced in paired-end mode on an AVITI system (Element Biosciences) by Quintara Biosciences (MA, USA).

FASTQ files were preprocessed with fastp. Read alignment and gene-level quantification were performed in Galaxy (usegalaxy.eu) using STAR in gene-counting mode against the mouse reference genome GRCm38/mm10. Gene-count outputs were imported into R for downstream analyses. Mouse Ensembl gene identifiers were mapped to gene symbols for pathway overlap and scoring.

Differential expression analysis was performed using DESeq2. Adjusted p values were calculated using the Benjamini–Hochberg procedure implemented in DESeq2, and results were summarized using an adjusted p value threshold of 0.10. DESeq2 independent filtering and outlier handling were applied according to the standard workflow (Love et al. 2014).

Pathway enrichment analysis was performed using Molecular Signatures Database (MSigDB) Hallmark gene sets (Liberzon et al. 2015) following the gene set enrichment analysis (GSEA) framework (Subramanian et al. 2005). To visualize sample-level pathway patterns, gene set variation analysis (GSVA) scores were calculated for OVX-altered Hallmark gene sets and displayed as row z-score heatmaps (Hänzelmann et al. 2013) (Appendix Figure 2).

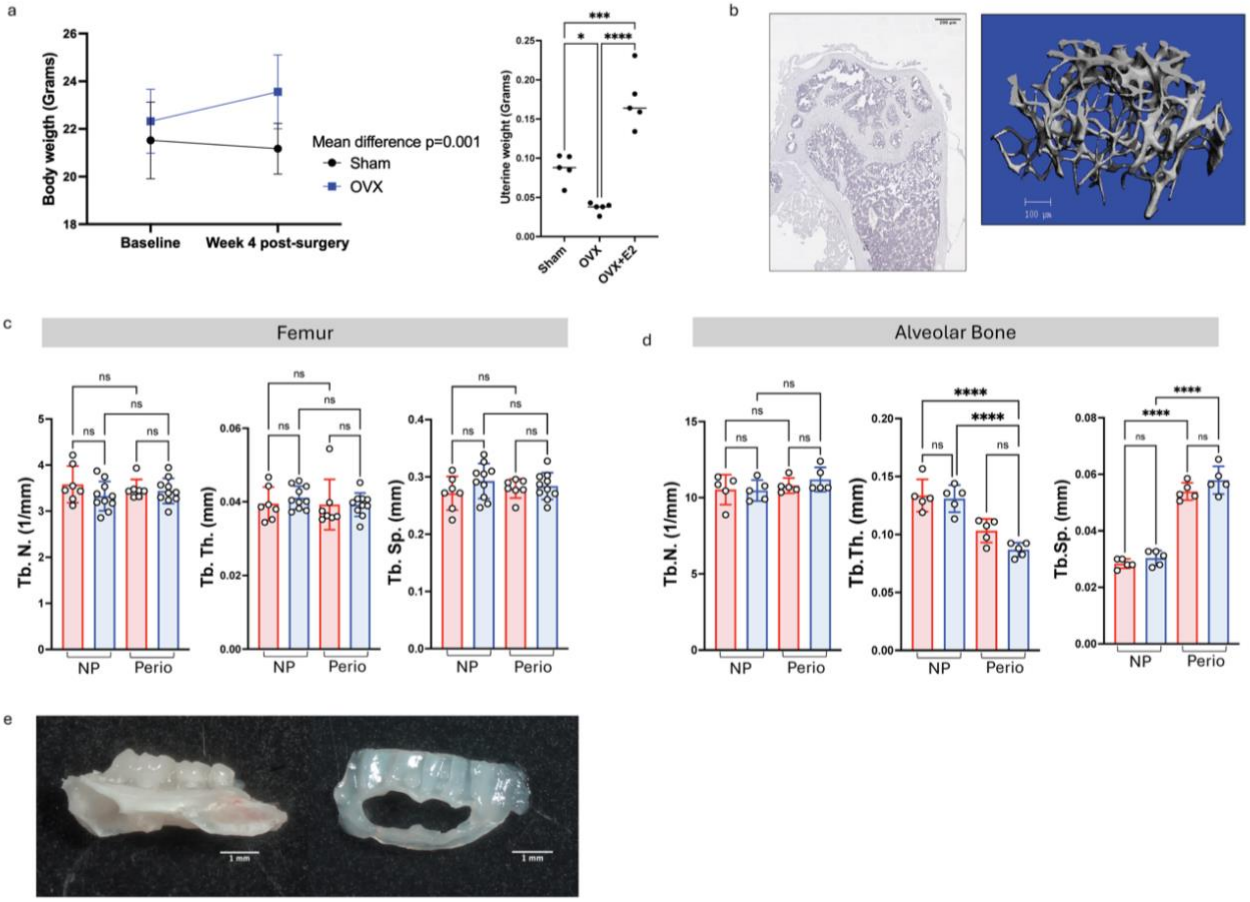


**Appendix Figure 1. a,** Ovariectomy efficiency confirmed by body-weight change in the longer timeline and by uterine weight in the shorter timeline. **b,** Representative osteocalcin IHC isotype-control image and distal femoral metaphysis micro-CT reconstruction. **c,** Femoral distal metaphysis trabecular number (Tb.N), trabecular thickness (Tb.Th), and trabecular separation (Tb.Sp) measured by micro-CT in the model shown in Figure 1a. **d,** Alveolar bone Tb.N, Tb.Th, and Tb.Sp measured by micro-CT in the model shown in Figure 1a. **e,** Representative mouse dentoalveolar complex and gingiva samples analyzed by bulk RNA sequencing. Sham, sham-operated; OVX, ovariectomized; OVX+E2, ovariectomized with estrogen replacement; Sham NP, sham-operated non-periodontitis; Sham Perio, sham-operated periodontitis; OVX NP, ovariectomized non-periodontitis; OVX Perio, ovariectomized periodontitis. Asterisks indicate statistical significance: $*p < 0.05$; $***p < 0.001$; $****p < 0.0001$.

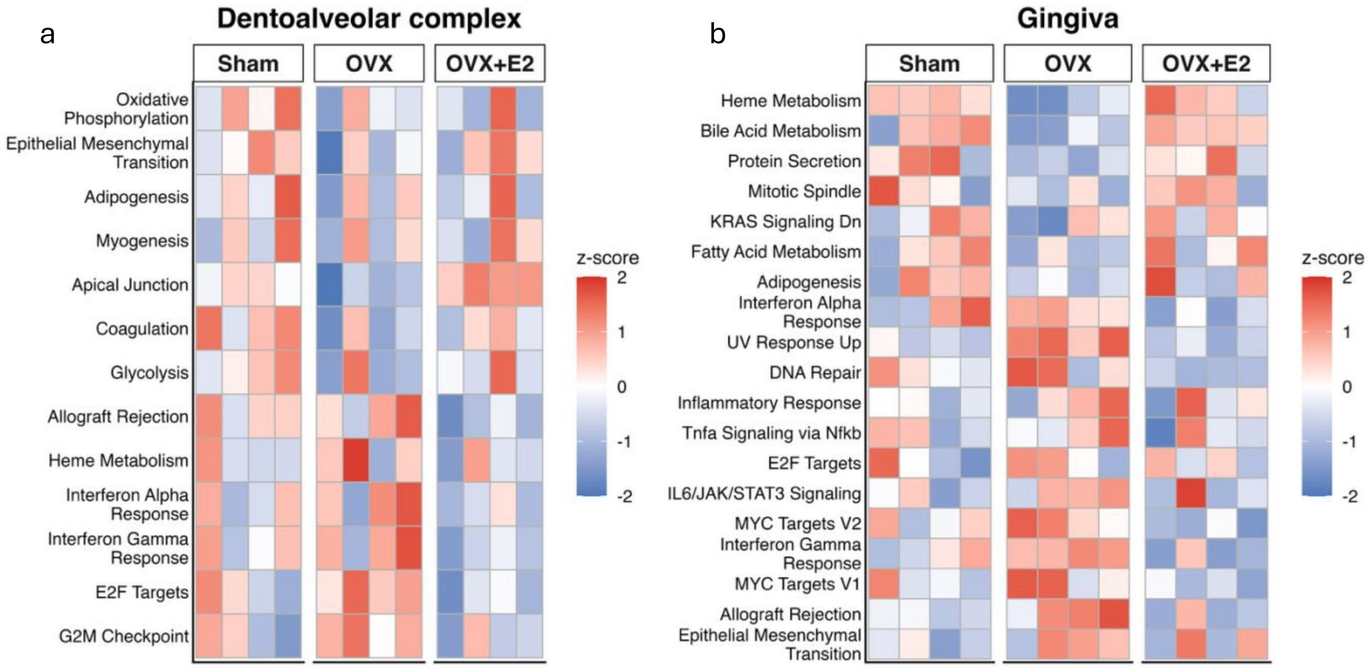


**Appendix Figure 2.** Gene set variation analysis (GSVA) heatmaps of OVX-altered, E2-responsive pathways. **a,** Dentoalveolar complex. **b,** Gingiva. Each row represents a Hallmark pathway that changed with OVX and moved away from the OVX pattern after E2 replacement. OVX-reduced pathways are placed at the top, and OVX-increased pathways are placed at the bottom. Sham, sham-operated periodontitis; OVX, ovariectomized periodontitis; OVX+E2, ovariectomized periodontitis with E2 replacement.
